# Phase-specific stimulation reveals consistent sinusoidal modulation of human corticospinal excitability along the oscillatory beta cycle

**DOI:** 10.1101/2023.04.25.538229

**Authors:** Marius Keute, Julian-Samuel Gebühr, Robert Guggenberger, Bettina Hanna Trunk, Alireza Gharabaghi

## Abstract

The responsiveness of neuronal populations to incoming information fluctuates. Retrospective analyses of randomly applied stimuli reveal a neural input-output relationship along the intrinsic oscillatory cycle. Prospectively harnessing this biological mechanism would necessitate frequency- and phase-specificity, intra- and inter-individual consistency, and instantaneous access to the oscillatory cycle.

We used a novel real-time approach to electroencephalography-triggered transcranial magnetic stimulation to precisely target 8 equidistant phases of the oscillatory cycle in the human motor cortex of male and female healthy participants. The phase-dependency of corticospinal excitability was investigated in ten different intrinsic frequencies (4, 8, 12, 16, 20, 24, 28, 32, 36, and 40Hz) and indexed by motor-evoked potentials (MEP) in the corresponding forearm muscle.

On both the individual and group level, we detected a consistent sinusoidal MEP modulation along the oscillatory cycle at 24Hz (χ^2^_2_ =9.2, p=.01), but not at any other target frequency (all χ^2^_2_ <5, all p>.08). Moreover, cross-validations showed also at 24Hz the highest consistency of the optimal phase between prospective (real-time) and retrospective (out-of-sample) testing (r=.605, p<.001), and across experimental sessions on three different days (r≥.45). The optimal corticospinal signal transmission was at the transition from the trough to the rising flank of the oscillatory 24Hz cycle.

Integrating real-time measurement and brain stimulation revealed that the sinusoidal input-output relationship of corticospinal signal transmission is frequency- and phase specific, and consistent within and across individuals and sessions. In future, this approach allows to selectively and repetitively target windows of increased responsiveness, and to thereby investigate potential cumulative effects on plasticity induction.

## Introduction

As a prominent feature of the brain, neural oscillations reflect synchronized periodic activity of neighboring neuronal populations (Buzsáki & Draguhn, 2004). In the context of neurophysiological measurements such as electroencephalography (EEG), these neural oscillations are often described as quasi-sinusoidal waves that can be spectrally characterized by power and phase (Cole & Voytek, 2017). According to the concept of communication-through-coherence, effective signal transmission is determined by the oscillatory phase of neuronal synchronization (Fries, 2015). In the human brain, this conceptual framework is studied by applying transcranial magnetic stimulation (TMS) pulses over the primary motor cortex during simultaneous EEG recordings. This enables us to determine the state of synchronized neuronal populations at the time of TMS while simultaneously capturing the induced stimulation effect indexed as the motor-evoked potential (MEP) amplitude in the corresponding muscle. When the response to TMS is indeed brain-state dependent, the induced MEP should follow the oscillatory pattern of the underlying brain dynamics at the time of perturbation.

Retrospective analysis of the MEP following randomly applied TMS pulses revealed less variable and more rapid corticospinal propagation at the optimum phase of beta frequency oscillations (13-30 Hz) (Torrecillos et al., 2020). Moreover, a beta frequency-specific gain modulation of cortical excitability was identified in both the resting and active motor system: Notably, both intrinsic (Khademi et al., 2018) and voluntarily modulated (Naros et al., 2020) beta power fluctuation influenced the cortical responsiveness to TMS, with high and low oscillatory activity reflecting inhibitory and excitatory states, respectively. Moreover, the motor cortex showed – independent of the oscillatory power fluctuations – a phase-dependent gain modulation along the oscillatory beta cycle that adhered to a unimodal pattern of increased responsiveness, and peaked along the rising phase of the oscillatory beta cycle with a diagonal shift of the maximum stimulation response with increasing frequency (Khademi et al., 2018).

Harnessing these findings for future interventions necessitates methodological and physiological preconditions: On the methodological side, phase-dependent modulation is critically dependent on precisely timing the stimuli to specific phases of the oscillatory cycle. However, previous interventions targeting phases of the oscillatory cycle required the application of prediction algorithms to compensate for latencies between measurement and stimulation, and were therefore imprecise. The higher the frequency, the more challenging the task of targeting a specific phase bin, since the stimulation timing needs to be faster than the cycle time of the target oscillation. However, previous EEG-triggered approaches hit the targeted negative and positive peaks of the alpha cycle (8-12 Hz) with standard deviations above 50° only (Zrenner et al., 2018, Madsen et al., 2019). This limit becomes particularly apparent when more precise phase targeting than hitting two opposite phases (e.g., peak vs. trough) in the alpha band is required to achieve the intended stimulation effect, e.g., by hitting a non-overlapping phase bin of 1/8 of the oscillatory beta cycle (Khademi et al., 2019). Importantly, a novel EEG-triggered TMS approach on the basis of integrated real-time measurement and stimulation achieved high precision, with standard deviations from 0.4° at 4 Hz to 4.3° at 40 Hz, thereby facilitating the selective targeting of at least 8 distinct phases across these frequencies (Guggenberger et al., 2023). Such precision would suffice to achieve the necessary phase specificity of TMS to maximize MEP increases in accordance with previous post-hoc estimations (Khademi et al., 2019).

On the physiological side, future phase-dependent interventions would necessitate robust frequency- and phase-specificity of the input-output relationship between stimulation and response, together with intra- and inter-individual consistency across sessions. Such evidence is, however, still lacking. We hypothesized that the integrated, real-time measurement and stimulation approach, which is now available, will enable us to prospectively confirm previous post-hoc observations, e.g., by identifying (i) a frequency- and phase-specific sinusoidal MEP modulation along the oscillatory cycle on both the individual and group level, (ii) a consistency of the optimal phase between prospective (real-time) and retrospective (out-of-sample) testing, and (iii) robustness across experimental sessions on three different days.

## Methods

### Design

Using a within-subjects design, we conducted a double-blind, randomized experimental study. In each experimental session, subjects received 1600 TMS pulses over the primary motor cortex with a uniformly jittered inter-stimulus interval (ISI) between 3.5 and 4.5 s. The primary outcome was muscle response to TMS (MEP). The exact timing of TMS pulses was controlled by the oscillatory phase of an EEG signal that was measured and analyzed in real time. We used ten target frequencies, ranging from the theta to gamma band (4 Hz to 40 Hz in steps of 4 Hz) and eight target phases (0° to 315° in steps of 45°). This resulted in 80 different frequency-phase combinations that were pseudo-randomized across stimuli (20 stimuli per combination). In addition, to keep subjects alert, 400 trials of a psychomotor vigilance task (PVT) were pseudo-randomly interspersed between TMS pulses, with the same ISI distribution.

### Participants

Twenty-one healthy adults (ten female) participated in our study. The mean age was 29.8 ± 9.7 years (range 21 to 58). All subjects were right-handed, reported no regular medical or recreational drug intake, and had no history of neurological, neurosurgical, or psychiatric treatment. An assessment of the inclusion and exclusion criteria was carried out via questionnaire before the intervention. The study protocol conformed to the Declaration of Helsinki and was approved by the Ethics Committee of the Medical Faculty at the University of Tuebingen. The study adhered to the current safety guidelines for application of TMS (Rossi et al., 2021), and none of the participants reported side-effects. All subjects gave their written informed consent prior to the experiment.

Each subject participated in three identical sessions that were scheduled at least 48 hours apart, but around the same time of day. Due to technical flaws, we lost data from four sessions (from four different subjects), resulting in a total of 59 sessions available for analysis.

### Apparatus

Electroencephalograpy (EEG) was recorded from 64 scalp channels using a BrainAmp amplifier (BrainProducts GmbH, Gilching, Germany) at a sampling rate of 1000 Hz and 16bit resolution. We used a TMS-compatible cap that adheres to the international 10-5 system (Easycap GmbH, Herrsching, Germany, online reference channel: FCz). Electrode impedances were kept below 5 kΩ. Electromyography (EMG) and electrocardiogram (ECG) were recorded from adhesive electrodes with Ag/AgCl contacts (Neuroline 720, Ambu A/S, Ballerup, Denmark). ECG was recorded in two-lead chest montage, EMG in bipolar belly-belly montage from the extensor digitorum communis (EDC), extensor carpi radialis (ECR), flexor carpi radialis (FCR), and flexor digitorum superficialis (FDS) muscles of both arms. Only data from the EDC, i.e., the target muscle for the experiment, will be reported here. In addition, we recorded breathing movements from a respiration belt (Vernier Software and Technology, Beaverton, USA; data not reported here).

For TMS, we used a triggerable MagPro X100 stimulator (MagVenture GmbH, Germany) with a figure-8 coil in combination with frameless stereotactic navigation (Localite GmbH, Germany).

TMS was triggered by a custom-built Integrated Measurement and Stimulation Device (IMSD; Neuroconn GmbH, Ilmenau, Germany). The IMSD provides a platform for categorizing brain states and controlling stimulation through real-time EEG analysis. During the experiment, the IMSD continuously measured EEG data from one bipolar channel (1 cm anterior and 1 cm posterior from C3) and extracted oscillatory phase at the target frequency across a 250 ms window by Discrete Fourier Transform (DFT) using the Goertzel algorithm (Sorensen et al., 1988) (Fig. 1). A target frequency and phase were pseudo-randomly picked for each trial from the set of predefined target frequencies and phases, and TMS was triggered by a 5 V TTL pulse once the EEG signal passed the target phase at the target frequency. Bipolar channel data from the IMSD were stored for offline analysis. Phase targeting with this real-time setup is highly accurate and precise, with mean deviations between the target phase and the actual triggered phase of 0.4° at 4 Hz to 4.3° at 40 Hz, and is described in further detail elsewhere (Guggenberger et al., 2023).

**Fig. 1.**
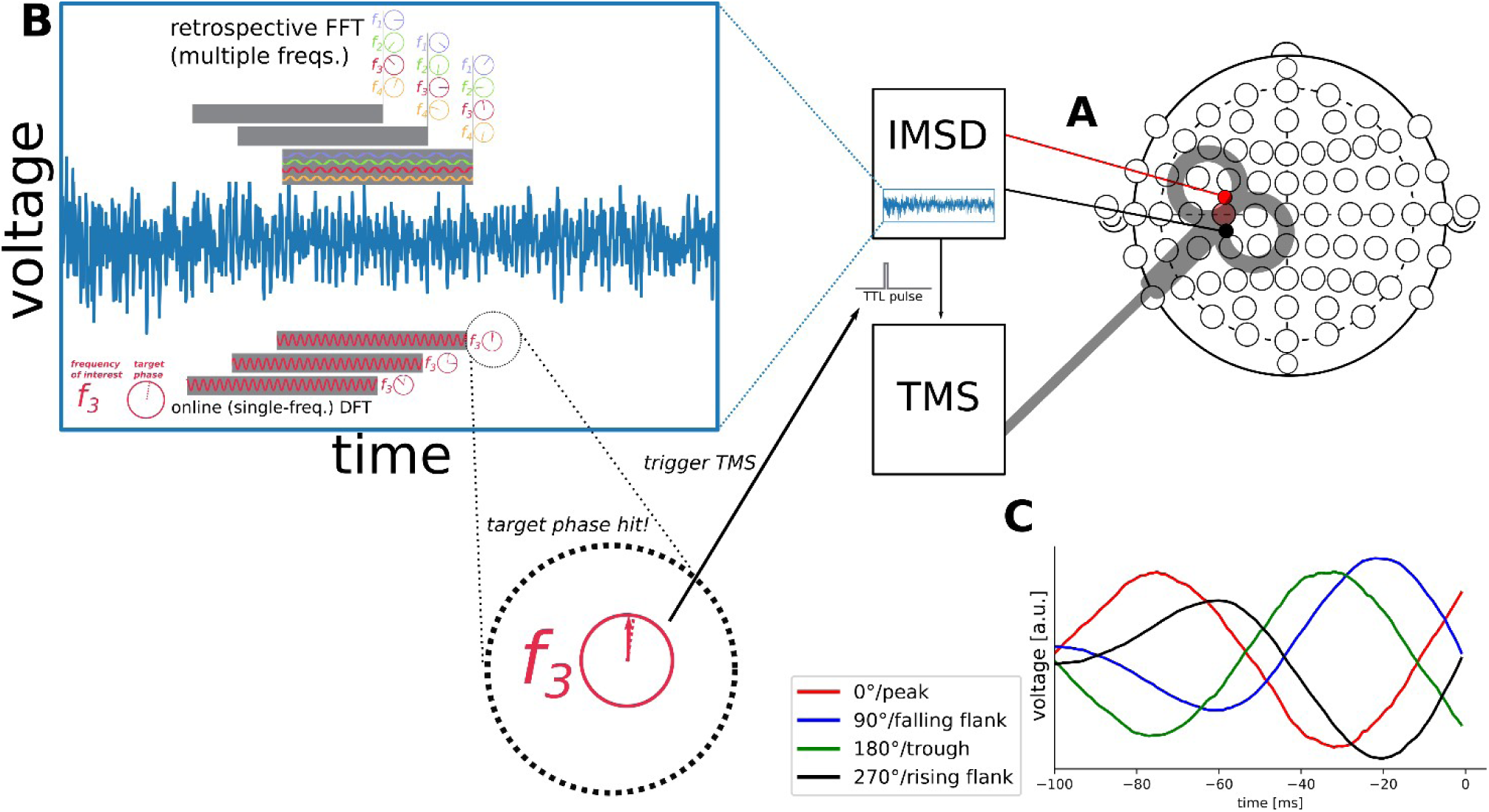
Experiment design and data analysis. A: EEG data was continuously recorded from 64 scalp electrodes (10-5 system), and from an additional bipolar channel 1 cm anterior (+) and 1 cm posterior (-) to C3. Data from this bipolar channel were analyzed in real time (∼ millisecond resolution) by a custom-built integrated measurement and stimulation device (IMSD). B: For each trial, a target frequency (between 4 and 40 Hz in steps of 4 Hz) and target phase (between 0 and 315 degrees in steps of 45 degrees) were chosen pseudo-randomly. The bipolar signal was analyzed online using a single-frequency (Goertzel) DFT on 250 ms time windows, and a 5V TTL pulse was sent to the TMS machine when the measured signal passed the target phase at the target frequency. Retrospectively, we calculated phases in the target and in all off-target frequencies from the bipolar IMSD signal using FFT with the same parameters as the online DFT. C: Grand average IMSD bipolar data traces from a 100 ms pre-pulse window are shown (right edge of plot is the trigger time). The traces show bandpass-filtered (8-16 Hz) grand median epochs from the 12 Hz target frequency trials at four different target phases (red: peak, blue: falling flank, green: trough, black: rising flank).

### Psychomotor Vigilance Task

Throughout the experiment, a white fixation cross was shown on the screen against a black background. Subjects were instructed to keep their gaze firmly on it. In PVT trials, the fixation cross disappeared and a white ‘X’ appeared on the screen for three frames (∼50 ms). Subjects were instructed to press a key on a PC keyboard with their left index finger as quickly as possible, and reaction time in ms was fed back visually.

### Hotspot search

The EDC hotspot was defined as the TMS coil position at which the highest MEP amplitudes (for details on MEP amplitude calculation see Section ‘Motor Evoked Potentials’) at the right EDC could be achieved. The search was initiated with a mapping of the left hemisphere using 40 TMS pulses at 45% of Maximum Stimulator Output (MSO). The stimulation intensity was increased in case no MEP was elicited. The coil was held tangentially to the top of the head and was rotated 45° relative to the parasagittal plane to achieve the optimal current flow for inducing the highest MEP amplitudes (Mills et al., 1992). The coil positions of the three highest MEPs were stimulated a further three times, and the location with the highest mean MEP amplitude was saved as hotspot, and used for all subsequent TMS pulses.

### Resting Motor Threshold

The RMT was defined as the lowest intensity at which 10 x stimulation at the EDC hotspot would result in MEP amplitudes greater 50 μV for at least half of the stimuli.

### Procedure

Subjects were seated comfortably in a reclining chair. Physiological measurements were prepared. The EDC hotspot and RMT were determined using the procedures described above. Six minutes of resting data were collected, alternating 30 s epochs with eyes open and eyes closed, respectively. During the subsequent intervention, the participants were requested to keep their right hand completely relaxed, while passively sitting and fixing their eyes on a fixation cross on the screen. Every 3.5 to 4.5 s (jittered), they either received a TMS stimulus or had to respond to a PVT trial. Stimuli were applied in 10 blocks of 200 stimuli (160 TMS pulses and 40 PVT trials per block), with a short (subject-terminated) break between blocks.

TMS pulses were applied at 120% of the RMT, using biphasic pulses with a pulse width of 280 μs. Since the intensity was set to 120% of the RMT, we expected to see MEP responses (> 50 μV) in practically every trial (Torrecillos et al., 2020).

### Motor Evoked Potentials

We determined MEP amplitude as the peak-to-peak range of the EMG signal over the EDC muscle in a time window of 15-60 ms after each TMS pulse. MEP amplitudes lower than 50 μV were excluded from further analyses. Furthermore, we excluded all trials with MEP amplitude > 5mV, all trials with muscle preactivation (MEP amplitude lower than three standard deviations of the MEP signal during the 500 ms preceding the pulse), and all trials with technical flaws in the IMSD (e.g., no signal available for assessment of phase adherence). A total of 19.2% of all trials were excluded on the basis of these criteria.

### Offline EEG analysis

For confirmatory retrospective analyses, we calculated the oscillatory phase of the bipolar EEG signal for each TMS pulse at all ten target frequencies, using Fast Fourier Transformation over the same time window as the real-time algorithm (250 ms, ending before the TMS trigger).

### Statistical analysis

#### Testing for time-related effects

MEP amplitudes were log transformed to approximate a normal distribution (Figure 2b). We used linear mixed-effects regression models (LMM) with per-session random intercepts to test, separately for each target frequency, for an effect of block number, pulse number within block (i.e., time into experiment and time since last break), and inter-pulse interval (i.e., time elapsed since last pulse) on log-MEP amplitude (Karabanov et al., 2021; Zarkowski et al., 2006; Zrenner et al., 2018). We tested for significance of predictors based on coefficient t-values, with Satterthwaite approximations for degrees of freedom.

**Fig. 2.**
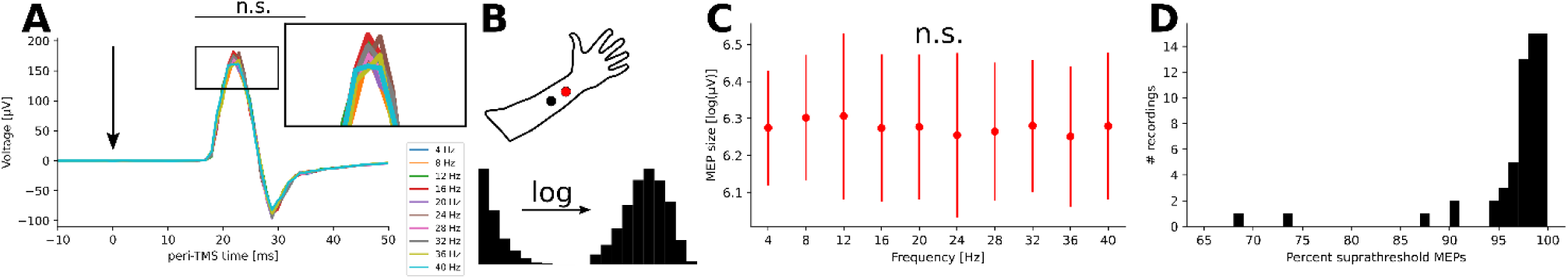
MEP overview. A: Session-wise percentage of trials with supra-threshold (>50μV) MEPs between all trials. B: Grand median EMG traces around TMS pulses (vertical arrow) per target frequency. Between 15 and 60 ms post-pulse, none of the time points showed a significant difference between frequencies (Kruskal-Wallis test, all p > .16), i.e., the overall EMG response did not differ between target frequencies. C: The approximate position of the EMG electrodes over the EDC is shown, as well as the distribution of extracted MEP amplitudes before and after log-transform. D: Log-MEP amplitudes per target frequency (mean and bootstrapped standard error across sessions). The values did not differ significantly between frequencies (Kruskal-Wallis test, p = .419).

#### Testing for phase-related effects

Next, on the basis of previous findings (Khademi et al., 2018; Torrecillos et al., 2020), we least-squares fitted cosine models to the log-MEP amplitudes. To render log-MEP amplitude and variance comparable between sessions, we calculated z-scores per subject, session, and target frequency (normalized log-MEP). The model was *M_i_ = a*cos(t_i_ + φ) + ε_i_*, with *M_i_* being normalized log-MEP amplitude for the i^th^ trial, *t_i_* being the corresponding target phase, *ε_i_*being the error term, *a* being the amplitude (which we will refer to as modulation depth) and *φ* the estimated optimal (i.e., producing the highest MEP amplitudes) phase shift, respectively (*φ=0 corresponding to a maximum at the positive peak of the oscillation*). No vertical offset was included in the model, since the data were z-scored (i.e., zero-mean). We created a null distribution for modulation depths by means of permutation, i.e., we randomly shuffled log-MEP values and target frequencies against target phases 1000 times, and fitted the model to the shuffled data.

We carried out cosine fitting and permutation testing both for pooled data from all sessions (i.e., assuming common model coefficients), and for each session individually (i.e., allowing model coefficients, in particular optimal phase, to differ between subjects / sessions). We expected higher model coefficients for individual models, since variations in optimal phase might cancel each other out on the group level.

The *group cosine fit* was considered significant if the modulation depth was greater than the 95^th^ percentile of the surrogate distribution. To ensure that results were not influenced by biases in the cosine model fitting process, we additionally confirmed the group-level cosine fit results by means of LMM, using the sine and cosine of the target phase as predictors for normalized log-MEP size, as recommended in the literature (Zoefel et al., 2019). We compared those models to null models using log-likelihood ratio tests (Model formula: logMEP ∼ cos(targetphase) + sin(targetphase) + (1|session); Null model formula:

logMEP ∼ 1 + (1|session); models with more complex random effects structures did not converge (Barr et al., 2013)).

The *individual cosine fits* were considered significant on the group level if the group mean of individual modulation depths was significantly higher than the group mean of the individual bias amplitudes (i.e., the average modulation depth achieved with permuted data), as determined by non-parametric Wilcoxon signed rank tests.

To assess phase consistency between individual cosine models, we calculated phase locking values (PLV) across the individual model estimates for optimal phase, using: 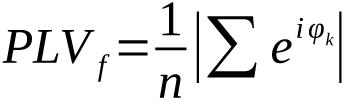 with f being target frequency, PLV_f_ being the PLV for the models for this target frequency, and φ_k_ being the estimated optimal phase in the k^th^ out of n individual models for this target frequency.

#### Testing stability of phase-related effects

To assess modulation stability within one session, we fitted cosine models to the offline calculated phases of all target frequencies for all trials, and compared optimal phase estimates with the real-time phase modulation by means of circular-circular correlation coefficients, as implemented in the pycircstat (www.github.com/circstat/pycircstat) toolbox (Berens, 2009; Jammalamadaka & Sengupta, 2001). Likewise, to assess within-session stability, we compared individual phase estimates obtained from the first and second half of each experimental session, respectively, to phase estimates for the whole session; and we compared optimal (online) phase estimates between sessions to assess intraindividual between-session stability.

We verified our analysis pipeline by visually inspecting a random selection of ∼1000 trials, together with extracted amplitudes and phases, and of ∼10% of the cosine fitting results, together with the underlying data.

## Results

### Motor-Evoked Potentials (MEP)

We first established that the measures were interpretable without systematic differences within and between sessions. Specifically, we confirmed that there were reproducible TMS responses with a high proportion (>85% of trials for all but two sessions, Fig. 2D) of supra-threshold MEP (>50 μV). One session with a low supra-threshold proportion (∼62%) was excluded from subsequent analyses due to a low number (n=224) of residual trials following artifact rejection.

Furthermore, average TMS-locked EMG traces (Fig. 2A, Kruskal-Wallis test, all p > .169) and log-MEP amplitudes (Fig. 2C, Kruskal-Wallis test, p = .419) were stable across different target frequencies, indicating that the phase-dependent modulations within each frequency could be compared between frequencies.

Notably, MEP amplitudes showed cumulative effects over time, i.e., they increased with block number (significant positive predictor at all frequencies, all t > 4.5, all p <.001) and with trial number within block (significant positive predictor at all frequencies except 12 Hz, all t > 2, all p <.04). The inter-pulse interval was a significant MEP predictor at 12 Hz (t = 2.5, p = .012) and 24 Hz (t = 2.9, p = .004).

### Sinusoidal modulation of MEP amplitude is frequency specific

Normalized MEP amplitudes per target frequency and target phase are shown in Fig. 3A. On the group level, we found a significant cosine modulation at 24 Hz in the permutation test (p = .027, Fig. 3B). Using linear mixed-effects regression models, we confirmed that phase had a significant effect at 24 Hz (χ^2^ = 9.2, p = .010, Fig. 3C), but not at any other target frequency (all χ^2^ < 5, all p > .08). The optimal phase occurred at the transition from the trough to the rising flank of the oscillatory 24 Hz cycle (Fig. 3A).

**Fig. 3.**
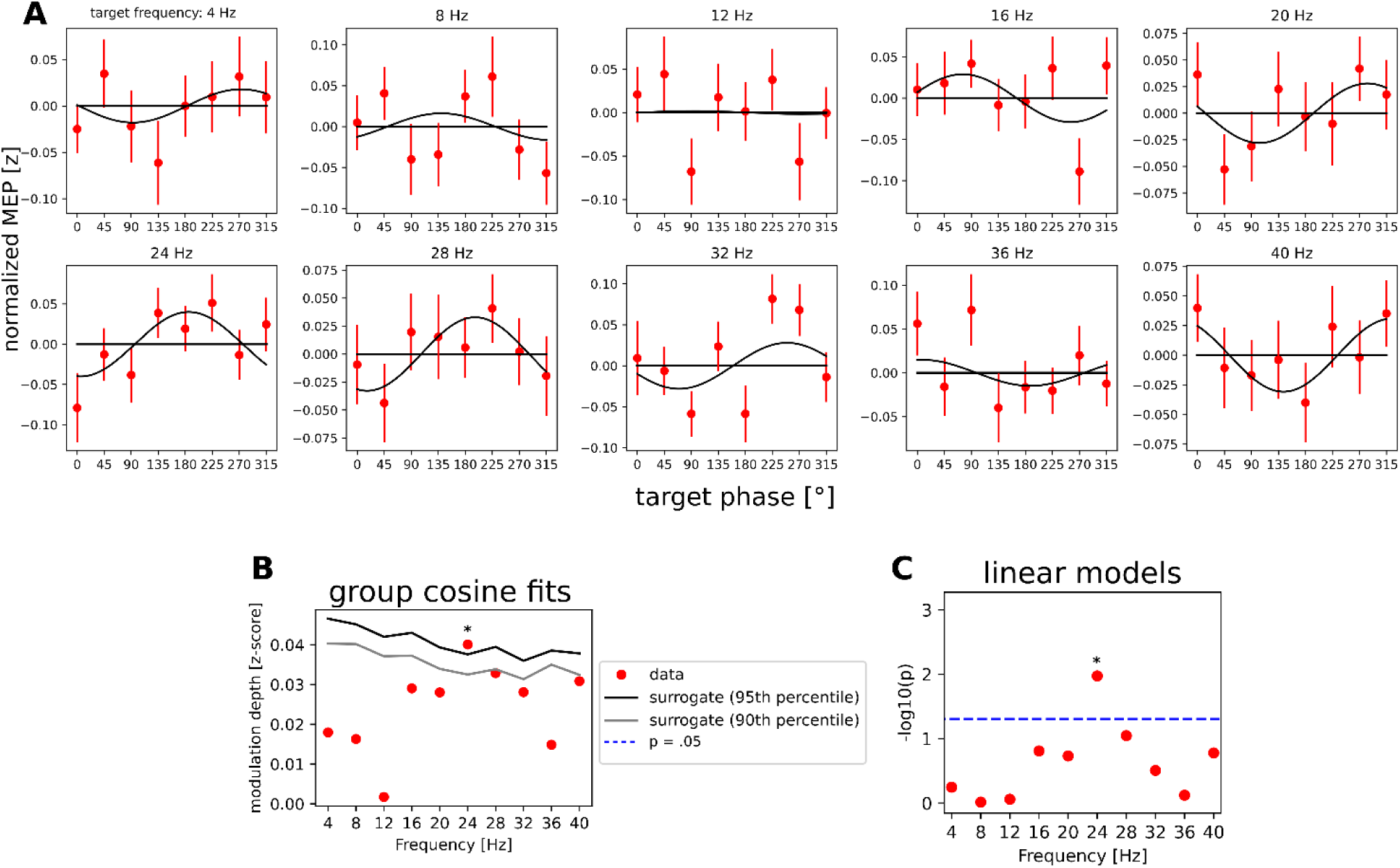
MEP group-level phase dependency. A: Normalized MEP amplitude (mean and bootstrapped standard error) across target phases for all target frequencies. Note that each subplot represents a non-overlapping set of trials (of ∼150 trials per session). The best-fitting group-level cosine is shown in black, corresponding to the modulation depths shown in Panel B: Cosine models. Group-level modulation depths (red), 95^th^ (black) and 90^th^ (gray) percentile of the surrogate distribution. A significant fit (p = .030) was found at 24 Hz. C: P-values (-log_10_(p)) from LMM regression between MEP amplitude and phase on the group level.

Controlling for the above-mentioned time-related predictors (block number, pulse number within block, and interpulse interval), i.e., by using regression residuals to test for MEP phase dependence) did not alter the overall pattern of MEP amplitudes across target frequencies and phases, leading to a consistent group-level effect of target phase on log-MEP amplitude at 24 Hz (χ^2^_2_ = 7.9, p = .019), but not at any other target frequency (all χ^2^_2_ < 5.9, all p > .05).

For individual models (Fig. 4A), we observed that cosine modulation depths were significantly higher than the mean surrogate (permutation) amplitudes at target frequencies 8 Hz (p = .003), 20 Hz (p = .008), 24 Hz (p = .003), 28 Hz (p = .033), 32 Hz (p < .001), and 40 Hz (p = .028). For most frequencies, the average optimal phase estimate of individual cosine models approximated the parameters of the group models (Fig. 4 B). However, individual models showed a wide distribution of estimated phase offsets, with the highest phase-specific consistencies (PLV > .18) at 24 Hz and 28 Hz.

**Fig. 4.**
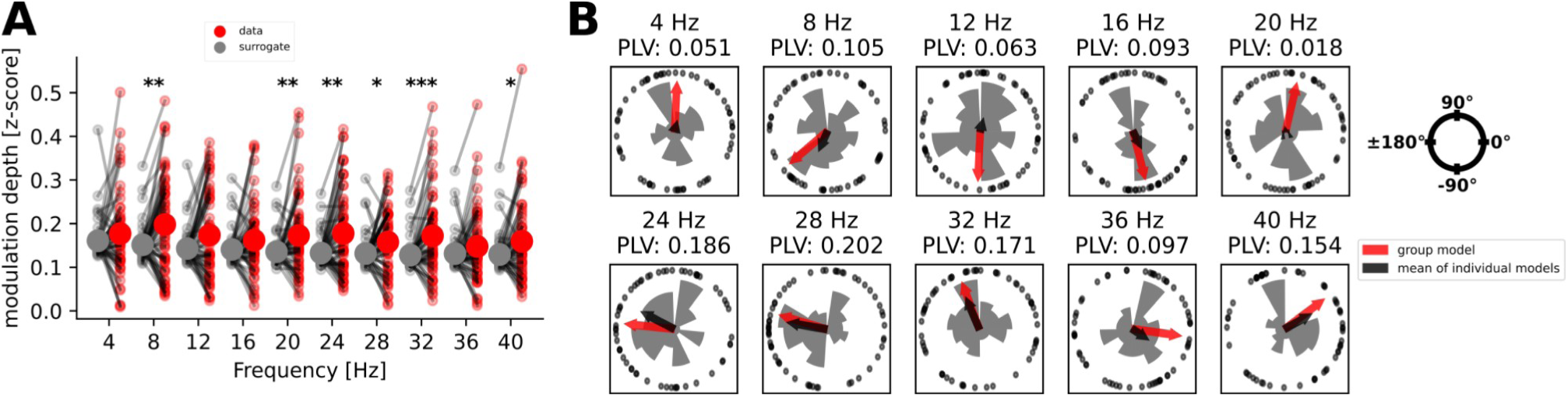
Individual MEP phase-dependency. A: Individual modulation depths and individual surrogate modulation depths at each target frequency (big dots: group mean). Individual cosine modulation is significantly greater than surrogate at 8, 20, 24, 28, 32, and 40 Hz. B: Individual values (black dots) and mean values (black arrows) for optimal phase estimates from individual cosine models and from group cosine models (red arrows). The estimated optimal phase is represented by the angle of the arrows. The length of the black arrows represents the average vector length of the mean phase and is proportional to the phase locking value (PLV). The length of the red arrows is arbitrary (set at 0.25).

### Intraindividual stability

We tested for stability of optimal phase estimates between real-time (prospective) and post-hoc (retrospective) models by means of circular correlation for all frequencies with individual phase-dependent MEP modulation (8 Hz, 20 Hz, 24 Hz, 28 Hz, 32 Hz, and 40 Hz). We found significant positive correlations between optimal phase estimates at 24 Hz (r = .605, p < .001), and, to a lesser degree, for 28 Hz (r = .342, p = .004), and 40 Hz (r = .253, p = .027). The difference in estimated optimal phase was below 10° at all frequencies except for 20 Hz (51.4°) (Fig. 5 A).

**Fig. 5.**
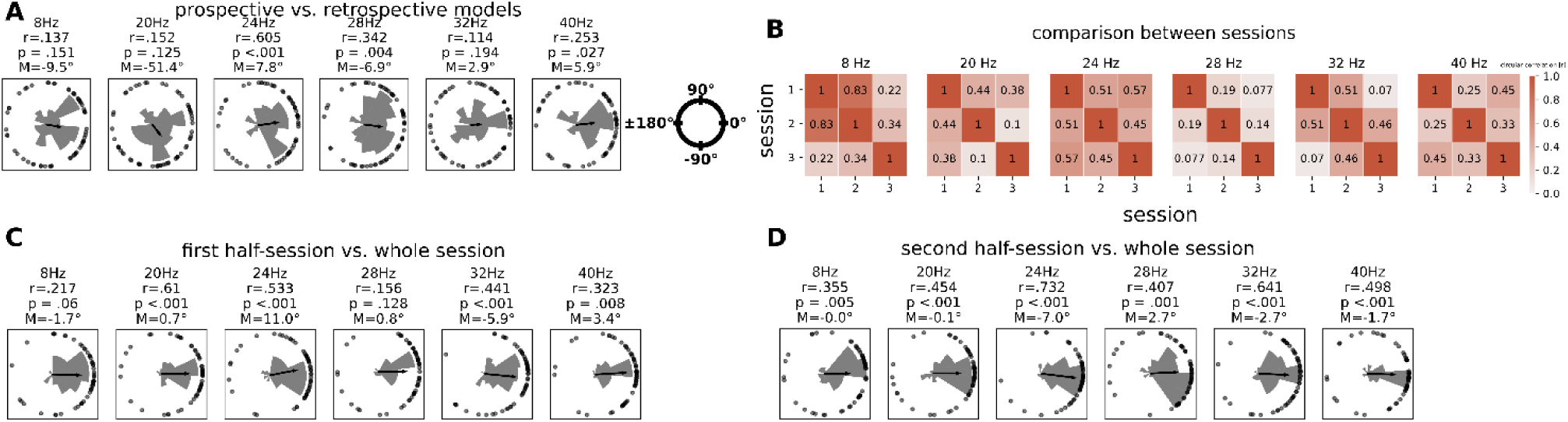
Stability of cosine modulation. A: Upper row: Scatterplots and correlations between individual real-time modulation depths and retrospective modulation depths (based on recalculated phases). B: Pairwise correlations (left panels) between modulation depths in the three experimental sessions, intra-class correlation coefficients (middle panel), and correlations between estimated optimal phases (right panel). C. Comparison between individual model coefficients based on the first half (∼800 trials) of sessions with the whole-session modulation depth. D: Same as C for second half.

The estimated optimal phases were consistently correlated across all three experimental sessions (r ≥ .45) for 24 Hz only (Fig. 5B).

In comparison to models estimated for the whole session, calculating cosine modulation depths for only the first or second half of the experimental sessions (∼800 trials per half-session), respectively, did not systematically alter optimal phase estimates (Fig. 5C/D). This indicates that cosine modulation does not critically depend on time-on-task.

## Discussion

In this study, we applied phase-dependent TMS with high precision and in real time across a range of target frequencies and phases. We aimed to establish whether corticospinal excitability (CSE), as quantified by MEP amplitude, was systematically modulated by the oscillatory phase of the EEG signal at the time of stimulation, as well as the frequencies at which such phase-dependent modulation occurred, and how stable this relationship was within and across sessions and participants. To this end, we modeled MEP amplitude as a sinusoidal function of 8 equidistant EEG phases at the time of stimulation for each of ten target frequencies between 4 Hz and 40 Hz in steps of 4 Hz on both the individual session level and group level.

By comparing CSE modulation in individual sessions against chance (randomly permuted phases), we established a significant CSE modulation for the whole upper beta band (20Hz, 24Hz, 28Hz, 32Hz), and for 8Hz and 40Hz. However, we found a group-level CSE modulation for 24Hz only, thus indicating interindividual consistency with respect to the optimal phase, i.e., the phase yielding the highest CSE at the time of stimulation.

Consistency of optimal phases was, furthermore, investigated by comparing the modulation depths between the phase-frequency combinations targeted in real-time (prospective model) and the post-hoc analyzed phase-frequency combinations that were simultaneously hit (retrospective model). Only for two frequencies in the upper beta-band (24Hz and 28Hz) were the optimal phases between prospective and retrospective models strongly correlated. Moreover, phase-dependency at 24Hz showed the highest intraindividual consistency across different sessions and measurement days. Taking all analyses into consideration, the strongest converging evidence for MEP phase-dependency was identified at a target frequency of 24Hz, with further – albeit less consistent – evidence for phase-dependency at other frequencies in the upper beta frequency band.

Previous work on EEG-triggered TMS revealed MEP differences when the negative vs. positive peak of the oscillatory alpha cycle was targeted in a group of participants selected on the basis of their high sensorimotor alpha power (Zrenner et al., 2018). However, this finding was not replicated when the same approach was applied in non-selected individuals (Madsen et al., 2019). Importantly, due to the temporal imprecision of these approaches (SD of ∼50°), a considerable number of stimuli would be outside the target phase for phase resolutions that are necessary for estimating sinusoidal modulation (Khademi et al., 2019; Guggenberger et al., 2023). Even though these methodological limitations have been overcome with novel integrated measurement and stimulation technology (Guggenberger et al., 2023) in the present study, further physiological challenges remain: Specifically, the neural sources generating periodic changes in CSE may have different positions and orientations relative to the scalp electrodes for different subjects, i.e., the peak and trough of the oscillations measured at the scalp electrodes may not be equivalent to the peak and trough of the oscillation at the underlying neural source. Along these lines, bipolar sensor-level recordings (as applied in this study) may differ from other EEG montages. Specifically, surface Laplacian spatial filtering might emphasize tangential dipoles in the cortex (Karabanov et al., 2021) more strongly than unprocessed sensor-level data (Buard et al., 2018; Hao et al., 2020).

In any case, for an interventional protocol to deliver repeated stimulation at the individual, maximally effective frequency and phase, it would be important that both can be estimated reliably and remain stable for at least the duration of the treatment. Our data revealed that such stable estimations are indeed possible. Moreover, we ascertained that modulation depths did not differ fundamentally between the early and late stage of the sessions. Even though we observed a relatively consistent phase-dependent CSE modulation for 24Hz across experimental sessions on three different days (r≥.45), our data do not suggest a strong trait-like effect.

In addition to phase-dependent modulations, we have shown that CSE also depends on phase-unrelated time factors, i.e., cumulative effects over time-on-task (Pellicciari et al., 2016). The inter-trial interval (ITI) between stimulation pulses, on the other hand, was not a consistent predictor of MEP amplitude as in previous studies (Julkunen et al., 2012), which might be related to the relatively small ITI variability in our study (jittered between 3.5 and 4.5s) (Schulz et al., 2014).

The present work may thus help to clarify previous contradictions in the literature concerning the CSE phase dependency as follows:

In our cross-validation experiments within and across sessions, and between individuals, we found no robust phase dependency in the alpha (mu) band. This is in line with recent work that applied Bayesian analyses and provided also evidence in favor of the null hypothesis (Karabanov et al., 2021). Converging evidence furthermore suggests that pericentral alpha (mu) activity is primarily located in the postcentral somatosensory cortex and may therefore influence CSE only indirectly (Karabanov et al., 2021, Zrenner et al. 2022). Moreover, the relation between stimulation intensity and the level of ongoing neural activity (Ogata et al., 2019), and the interaction between oscillatory phase and power may influence CSE to an extent unexplained by each of these features alone (Hussain et al., 2019). Also, individual post-hoc significance testing for a sinusoidal fit yielded alpha (mu) phase dependency in one third of individuals only (Zrenner et al., 2023). Furthermore, minor differences in the methodological choices may critically affect the sensitivity to detect the complex relationship between oscillatory activity and corticospinal excitability (Karabanov et al., 2021) and even lead to erroneous phase estimations (Khademi et al., 2023).

In contrast, this wok revealed robust and consistent sinusoidal CSE modulation along the oscillatory beta cycle. This is in line with previous work on the beta phase dependency of CSE using post-hoc analysis or transcranial alternating current stimulation (Guerra et al., 2016; Raco et al., 2016; Schilberg et al., 2018; Khademi et al., 2018; Naros et al., 2020; Torrecillos et al., 2020) and of the timing of voluntary movements (Hussein et al., 2022). Interestingly, a previous post-hoc analysis (Torrecillos et al., 2020) found a group-level sinusoidal MEP amplitude modulation between 23 and 25Hz only, which corresponds exactly with our findings. Of note, another online study suggests that the phases of alpha (mu) and beta oscillations have an opposite effect on corticospinal excitability (Wischnewski et al., 2022). However, this surprising observation needs further clarification; applying high-resolution phase sampling along the oscillatory cycle like in the present study may allow for robust modeling of input-output relationships and specific comparisons between different frequencies.

### Limitations and perspectives

The main challenge with *calculating* phases of brain oscillations is that choices of the phase extraction method (e.g., short-time Fourier Transform vs. instantaneous phase from the analytic signal), window length and tapers (where applicable) can have a profound impact on phase estimations (Zrenner et al., 2020, Khademi et al., 2023). The common FFT-based methods will return a phase estimate for any frequency (up to the Nyquist frequency) in any signal, regardless of whether or not an oscillation at the frequency of interest is actually present. The presence or absence of oscillatory activity can, in turn, be difficult to determine, particularly in a real-time setup. There are, however, algorithmic solutions to this problem, e.g., decomposing power spectra into periodic (oscillatory) and aperiodic parts by means of curve fitting (Donoghue et al., 2020). In addition to being distorted by aperiodic parts of the spectrum, phase estimates are highly sensitive to random noise. It is therefore crucial to eliminate as much noise as possible during the EEG measurement. As recently suggested (Wodeyar et al., 2021), future studies should consider more advanced online phase calculation methods such as Kalman filtering to account for noise.

The most difficult aspect of *interpreting* phase estimates, on the other hand, is that it is notoriously unclear as to what an oscillation measured outside the brain physiologically represents. Intracranially measured oscillations are usually interpreted in terms of depolarization or hyperpolarization of a local neuron population (Zanos et al., 2018), whereas EEG sensor-level data represent an unknown mixture of neural sources, and isolating one oscillation of interest from all background activity is often challenging and ambiguous. Moreover, it remains unclear as to whether sinusoidal oscillations really are the best way to represent rhythmic brain activity in the first place. However, sinusoidal signal representation is inevitable whenever phase extraction methods based on Fourier Transform are used. Brain activity has been shown to have prominent non-sinusoidal features (Cole & Voytek, 2017), and alternative phase extraction methods, e.g., empirical mode decomposition (Muñoz-Gutiérrez et al., 2018) or cycle-by-cycle analysis (Cole & Voytek, 2019), can potentially account for this to some extent.

In conclusion, integrating real-time measurement and brain stimulation revealed a sinusoidal input-output relationship of corticospinal excitability in the beta frequency band that was consistent within and across individuals and sessions. The optimal corticospinal signal transmission was frequency- and phase-specific at the transition from the trough to the rising flank of the oscillatory cycle. In future, this approach may allow to selectively and repetitively target windows of increased responsiveness, and to thereby investigate potential cumulative effects on plasticity induction or dysfunctional neural oscillations (Cagnan et al., 2017; Holt et al., 2019).

## Authorship contribution statement

Marius Keute: Data curation, Formal analysis, Writing - original draft, Visualization. Julian-Samuel Gebühr: Methodology, Software, Writing - review & editing. Robert Guggenberger: Methodology, Software, Writing - review & editing. Bettina Hanna Trunk: Investigation, Data curation, Writing - review & editing. Alireza Gharabaghi: Conceptualization, Writing - original draft, Supervision, Project administration, Funding acquisition.

## Acknowledgments

This work was supported by the German Federal Ministry of Education and Research [BMBF16SV8174, INERLINC]. We acknowledge support from the Open Access Publishing Fund of the University of Tuebingen.

## Data and code availability

The data that support the findings of this study are available for researchers from the first author upon reasonable request. Python code used for analysis will be made available on Github (https://github.com/mkeute/phase_dependent_MEP).

